# Characterizing expression of candidate genes of a glutamate receptor pathway in *Arabidopsis thaliana* using real-time RT-PCR

**DOI:** 10.1101/2020.01.24.918599

**Authors:** Julie Wurdeman, Tessa Durham Brooks

## Abstract

The *Arabidopsis thaliana* genome contains twenty genes that are analogous to mammalian ionotropic glutamate receptors. There are sixteen mammalian glutamate receptors, which are best known for their roles in neuroplasticity, learning, and memory. The large number of glutamate receptors in *A. thaliana* suggests they play important roles in the plant’s growth and development, possibly serving to regulate function like they do in non-excitable mammalian tissues. A specific glutamate receptor, *GLR3.3*, is highly expressed in root tissue of plants, and has been found to promote stronger, more coordinated curvature development during the process of gravitropism. Gravitropism is the ability of a plant to change its orientation to that of the gravity vector when displaced from its gravitational set point angle (GSPA). A previous association study identified six candidate genes which were correlated with the same phenotypic characteristics of gravitropism as *GLR3.3*. Utilizing real time RT-PCR (qRT-PCR) expression profiles were created for each candidate gene, including *GLR3.3.* A qRT-PCR method was developed to provide a more quantitative and sensitive way for measuring gene expression than traditional PCR methods. Furthermore, MIQE (Minimum Information for Publication of Quantitative Real-Time PCR Experiments) guidelines were followed to ensure data robustness. Expression profiles that were similar to *GLR3.3* were hypothesized to be good candidates as cell signaling components of this novel pathway. This is the beginning of a process that will identify a *GLR*-dependent pathway, the role of this novel pathway in the gravitropic response, and the influence of *GLRs* in plant physiology.

## Introduction

The *Arabidopsis thaliana* genome was one of the first genomes to be comprehensively sequenced, making it a valuable model for identifying and determining gene function (The Arabidopsis Genome Initiative 2000). The genome sequence revealed that *Arabidopsis* contains 600 genes that are hypothesized to encode for membrane transport proteins, of which 20 are similar to mammalian ionotropic glutamate receptors (iGluRs) (Lacombe et al. 2001). Glutamate receptors are ligand gated ion channels that bind amino acids. Activation of the receptor causes depolarization of the cell membrane through ion channels (Sliverthron 2013). Additionally, glutamate receptors interact with cell signaling pathways. iGluRs have been identified to play roles in synaptic plasticity, affecting processes such as learning and memory, the development of chronic pain and anxiety, and the manifestation of certain neurodegernative diseases including Alzheimers in mammals (Asztely and Gustafsson 1996; Bowie 2008). iGluRs have also been identified in a number of non-excitable mammalian tissues such as the heart, lungs, intestine, kidney, bones, and endocrine cells (Gill and Pulido 2001). In these non-nervous tissues, iGluRs are involved in regulation.

In A. *thaliana*, similarly related ion channel forming Glutamate Receptor-Like genes *(AtGLRs)* are categorized into three clades based on their amino acid binding properties (Lacombe et al. 2001). These receptors are hypothesized to be most similar to the extensively researched NMDA clade of iGluRs, because of the *AtGLRs’* similar function as nonselective cation channels that allow for calcium conductance (Cheffings 2001). However, it remains unclear why *A. thaliana*, and plants in general, have such a large number of glutamate receptors considering their lack of a central nervous system (Mayer and Armstrong 2004). It is hypothesized that glutamate receptors have a regulatory purpose in plants, in a way similar to that of non-excitable mammalian tissues (Davenport 2002). Clade three *AtGLRs* are highly expressed in root tissues, thus, making them the primary candidate for glutamate receptors involved in the process of root gravitropism (Chiu et al. 2002). In support of this idea, *GLR3.3* has been found to play a regulatory role in the process of root gravitropism.

Gravitropism has been a topic of study for many decades, however much is unknown about the subject and new mechanistic pathways continue to be discovered. Gravitropism allows plants to realign their organs with gravity and resume growth at the gravitational set point angle (GSPA), a defined angle from the gravity vector (Blancaflor and Masson 2003). Roots display this phenomena, with each species of plant defining a unique GSPA (Firn and Digby 1995). Specifically, in roots, the root’s tip detects a change in gravity induced by the environment (e.g. effects from wind or disturbance of plant from level ground). This activates a signaling cascade which induces the hormone auxin to be differentially distributed between the upper and lower surface of the root, which results in an altered growth rate (Evans 1991). When the process is complete the root tip is realigned with the GSPA. This process is complex and multiphasic. Recently, a specific *AtGLR* in clade three, *GLR3.3*, was discovered to be expressed midway through the gravitropic process (Miller et al. 2010).

When *A. thaliana* is rotated 90 degrees, the roots of the plant direct their growth toward the gravity vector. This process is divided into two phases: the first phase consists of rapid tip angle development during the first 30-40 degrees of tip angle accruement; the second phase consists of more graduate tip angle development to the set point angle (Durham Brooks et al. 2010). During this two phase process the root can overshoot its trajectory, but will correct its growth accordingly. Approximately 2-4 hours after a gravitropic stimulus is presented, when the root has bent by 30-40 degrees, mutants in *GLR3.3* show a deceleration and slower tip angle accruement than wild type seedlings.

*GLR3.3* is active during the second phase of the gravitropic response; this was determined by the abnormal phenotype of mutant *glr3.3* seedlings, which is observed during this second phase. Mutants of *GLR3.3* showed slower tip angle accruement and more stochastic tip angle development during the second phase of the response, suggesting that *GLR3.3* is responsible for promoting curvature and ensuring a uniform response (Miller et al. 2010). The transition from rapid to slower tip angle development during root gravitropism may represent a period when information is processed and adjustments are made. This proposed function is consistent with the role of a glutamate receptor in a regulatory process, as is seen in mammalian cells (Gill & Pulido 2001). This regulatory aspect of the mechanism of root gravitropism has not yet been characterized, and the cell signaling components have not yet been identified. Therefore, the goal of this project is to uncover additional components of this novel pathway.

A previous association study identified six possible candidate genes which correlated the phenotypic and genotypic characteristics of gravitropism (Merithew 2012). In this study, expression profiles were created for each of these genes to determine if their expression profile is similar to that of *GLR3.3*, and therefore, a candidate for the cell signaling pathway. These genes were identified based on their association with the phenotype at approximately the same time as the *GLR3.3* locus and annotated functions consistent with a role in cell signaling. The functions of these genes include transcription factors, ion channels, and kinases. In these previous experiments, *A. thaliana* root tissue was collected at 3.5, 4.5, and 5.5 hours after being gravitropically stimulated. RNA was extracted from this tissue and RT-PCR was performed to determine whether the candidate genes were expressed in the root with *GLR3.3*. It was found that all six of the candidate genes were expressed in the root with *GLR3.3* during the root gravitropic response, indicating that all candidate genes could be components of a *GLR3.3* signaling pathway. This result was very unexpected, and to confirm the validity of the results the candidate genes need to be tested with a more sensitive method. Instead of using RT-PCR, qRT-PCR will allow the relative quantification of levels of expression. Using information obtained through qRT-PCR and previously collected data, it is possible that components of the glutamate cell signaling pathway could be identified. Candidate genes that are similar to *GLR3.3* will show similar expression profiles, while dissimilar expression in candidate genes indicates less probable association to *GLR3.3.* This is the beginning of a process that will identify a *GLR*-dependent pathway, the role of this novel pathway in the gravitropic response, and the influence of *GLR*s in plant physiology.

## Materials and Methods

### Primer Design

Merithew (2011) found 6 genes highly correlated with the gravitropic response of *GLR3.3.* Using the published Minimum Information for Publication of Quantitative Real Time PCR Experiments (MIQE Guidelines) and guidelines provided by Bio-Rad, primers were designed for these 6 genes using NCBI’s Primer BLAST program (http://www.ncbi.nlm.nih.gov/). The parameters for the primers were a product size of 70 to 150 and a melting temperature of 50-60°C. It was preferred the primer span an exon-exon junction and the GC content be between 50-60% The divalent cation concentration was set at 1.2 and the dNTP concentration was 0.2. Preference was shown to primers that began with guanine and cytosine and ended with adenine and thymine. Primers were also analyzed to prevent unwanted annealing to non-targeted regions of the genome. IDT Technologies Oligoanalyzer was used to ensure no dimer or hairpin structures could be formed with the primer pairs. Primer sequences are displayed in Table 1.

**Table 1:**
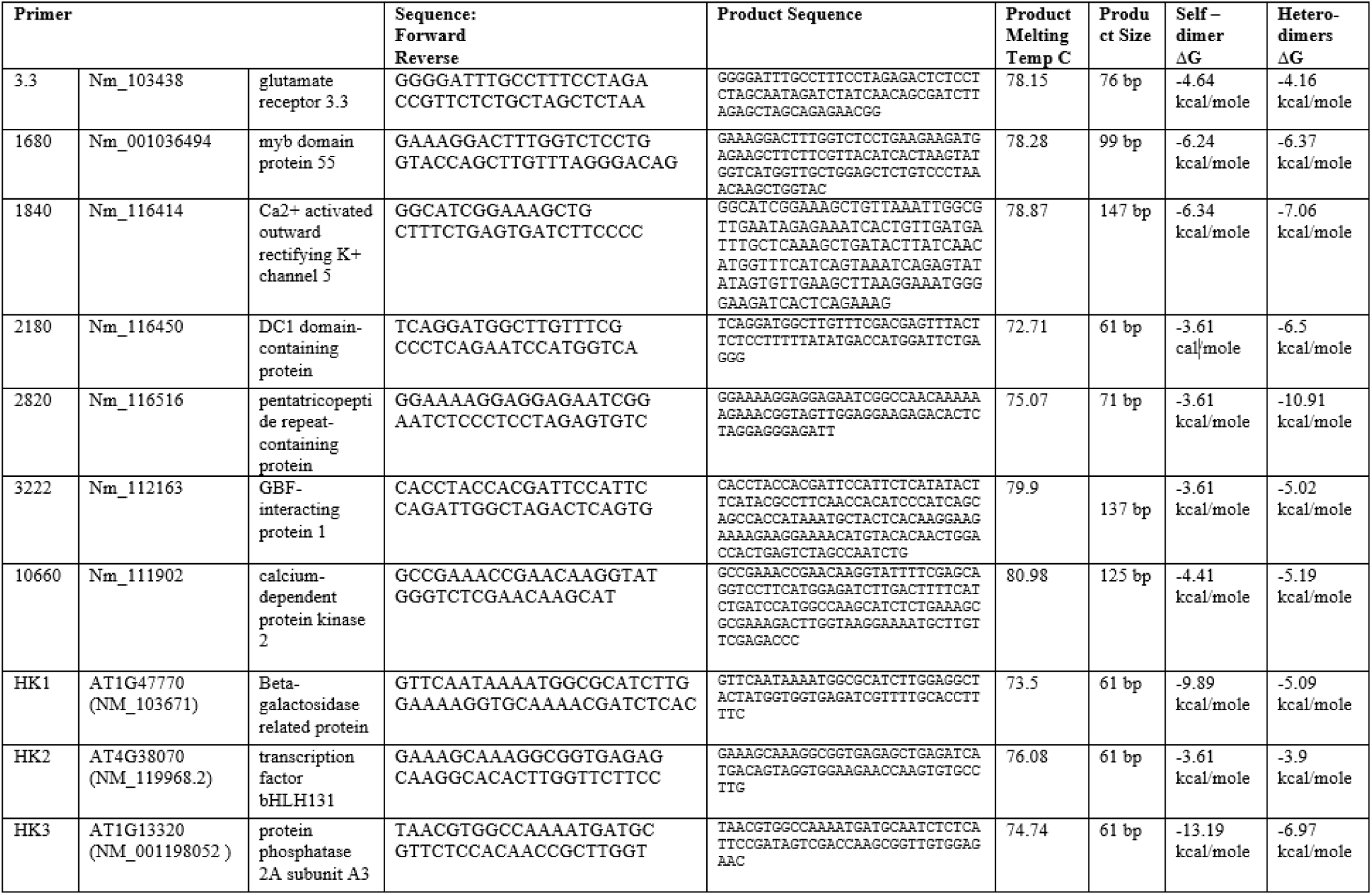
Primer Sequences.

### Planting

Tissue was collected using the *Arabidopsis thaliana* ecotypes, Cape Verde Islands (Cvi, 163) and Landsberg erecta (Ler, 164). Seeds from both ecotypes were planted in a standard 100 × 15 mm petri dish with 1% agar medium with minimal media containing 1 mM KCl, 1 mM CaCl_2_, 5 mM 2-[N-morpholino]-ethanesulfonic acid, and adjusted to a pH of 5.7 with 1,3-bis[tris(hydroxymethyl)methylamino] propane. Seeds were sterilized with a wash of 2% Triton X plus 70% ethanol followed by a wash 70% ethanol. Seeds were planted in 3 parallel rows with approximately 50 seeds per line, and a total of 150 seedlings per plate. A total of 12 plates were planted for each experimental replicate. Plates were incubated at a 4°C for 5 days. Plates were then placed in a contraption that ensured they remained vertical at all times and placed in a growth chamber delivering continuous white light at a fluency rate of 60.6 μmoles/(m^2^*sec) and a temperature of 25°C.for 3 days.

### Tissue Collection

After 3 full days of growth, plates were carefully transferred to a RNase free collection zone. Plates were placed in holders that ensured that they were kept vertical. Tools needed to collect tissue included an RNase free scalpel, forceps, pre-labeled microcentrifuge tubes and a sterile cooler filled with liquid nitrogen. All plates, excluding three control, non-gravistimulated 0 hour plates, were turned 90° to induce gravitropism. To collect tissue, plates were held carefully not to be rotated, and the roots of the seedlings were cut with the scalpel right below the root-shoot junction for an entire row on the plate. This row was then collected with a dragging motion of the forceps across the roots. It was important not to collect agar or to disturb the root excessively. Once a row of tissue was collected in the forceps, it was immediately flash frozen in the pre-labeled microcentrifuge tube. This process was then repeated for the second and third rows on the plate. Time points were collected at 0 hours, 3.5 hours, 4.5 hours, and 5.5 hours. The tissue of 3 plates were placed into a single microcentrifuge tube for each time point. Once the tissue was collected it was placed in a −70°C freezer.

### RNA Extraction

RNA extraction was performed with the collected tissue. The process occurred in a sterile RNase free zone. A single tube of tissue was ground using an electric drill and a plastic Rnase free pestle. While the sample was in liquid nitrogen, the pestle was inserted into the microcentrifuge tube containing the root tissue. Tissue was ground for three to four times for 45 seconds to one minute. After each grinding the tube was momentarily removed from the liquid nitrogen to tap the bottom of the tube to attempt to displace any compact pellet that had formed at the bottom of the tube, which could prevent complete grinding of the sample. After grinding the tissue into a fine white powder, a Qiagen RNeasy Plant mini kit was used to extract RNA. The “Purification of Total RNA from Plant Cells and Tissues and Filamentous Fungi” protocol provided with the kit was followed, including the optional DNAse Digestion steps. A total of 40 μL of RNA was collected from each tissue sample. RNA was analyzed on a NanoDrop 2000 Instrument to ensure purity of the sample.

### Standard Curve Construction

Real-Time PCR was completed using a Bio-Rad Thermocycler. All experimental RNA samples were reacted with Bio-Rad Iscript Reverse Transcription Supermix to create cDNA samples with the provided reaction protocol. At the same time a No Reverse Transcriptase control sample was created for each RNA sample. Using information from the nanodrop values, 100ng/μl was used per reaction. All primer pairs were tested for their optimal melting temp by utilizing the temperature gradient setting on the Bio-Rad thermocycler. The highest and lowest values were found and the average was calculated. This average was input into the Gradient Calculator in the CFX manager software to calculate a range of 65-45°C. The optimum temperature for all primers was found to be 57.6°C.

Initially, all primers were validated by creating a standard curve. To do this an available cDNA sample was serially diluted on a range from 8ul to 8 × 10^−7^ μL per reaction in 8 10-fold dilutions. When performing the serial dilution it was very important that volumes pipetted were accurate. If concentrations of the cDNA vary, the outcome is very evident in high efficiency values for the standard curve. SsoFast EvaGreen Supermix was used for a reaction volume of 20μL per sample. All reactions were technically repeated in triplicate. A mastermix was made for each dilution using 10 μL of supermix, 0.04 μM forward primer, 0.04 μM reverse primer, cDNA, and RNase free water varying, and bringing the volume to 20μL. The thermocycler conditions are shown in Figure 1. An additional melt curve from 70-90°C was performed after amplification in order to determine if multiple products were present in the samples. Standard Curves were considered acceptable if they showed at least 4 dilutions from the series, had an R^2^ > 0.98 and an efficiency value between 95%-105%.

**Figure 1:**
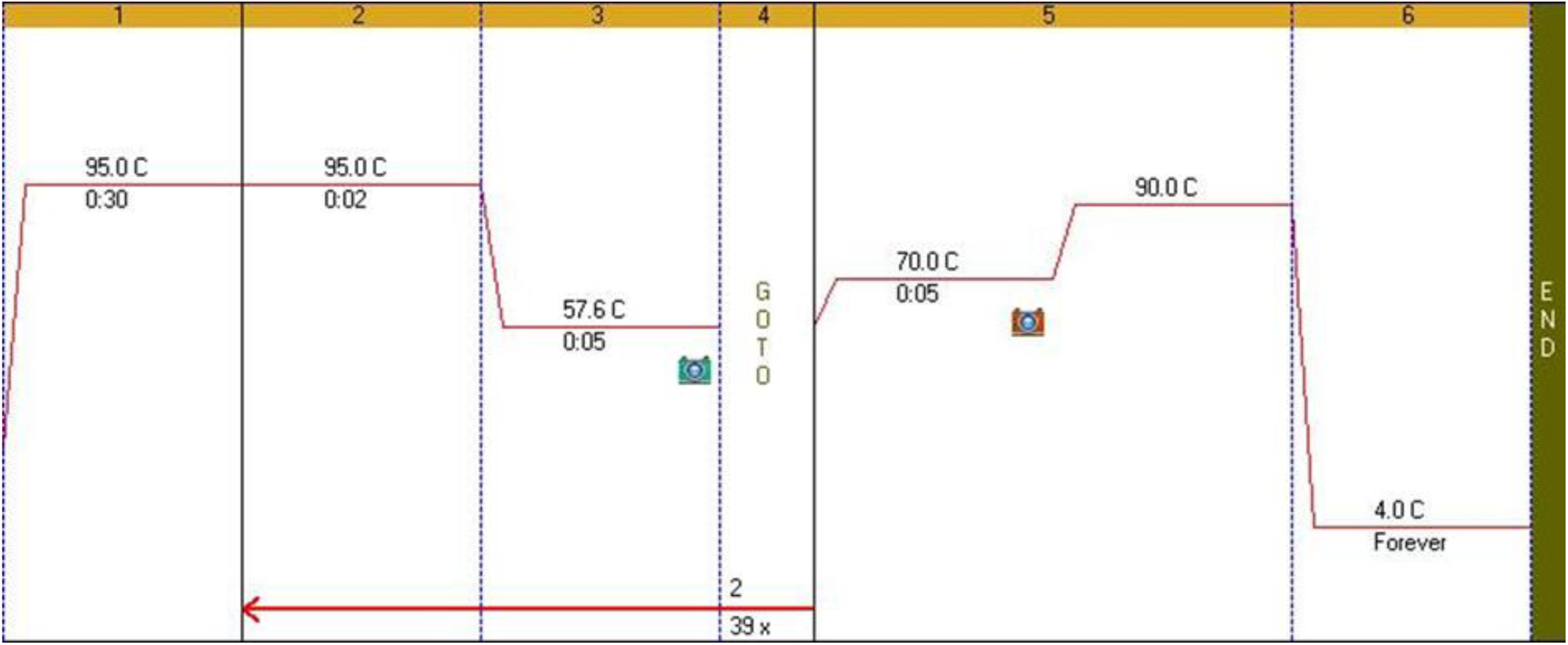
Thermocycling conditions shown as programmed in the Bio-Rad CFX software program.

### Gene Study

Once primers were validated, a gene expression study was performed using the same supermix and thermocycling conditions as was used in the standard curves. The reactions differed in that a constant cDNA volume of 0.05μL was added to each reaction. This amount was selected by analyzing all primers’ standard curve data and finding the median concentration on each curve. Data was analyzed using the CFX Manager Software. The plate setup used to run a gene study is shown in Figure 2. See *Appendix A* for more detailed instructions of how to use and program the CFX manager software.

**Figure 2:**
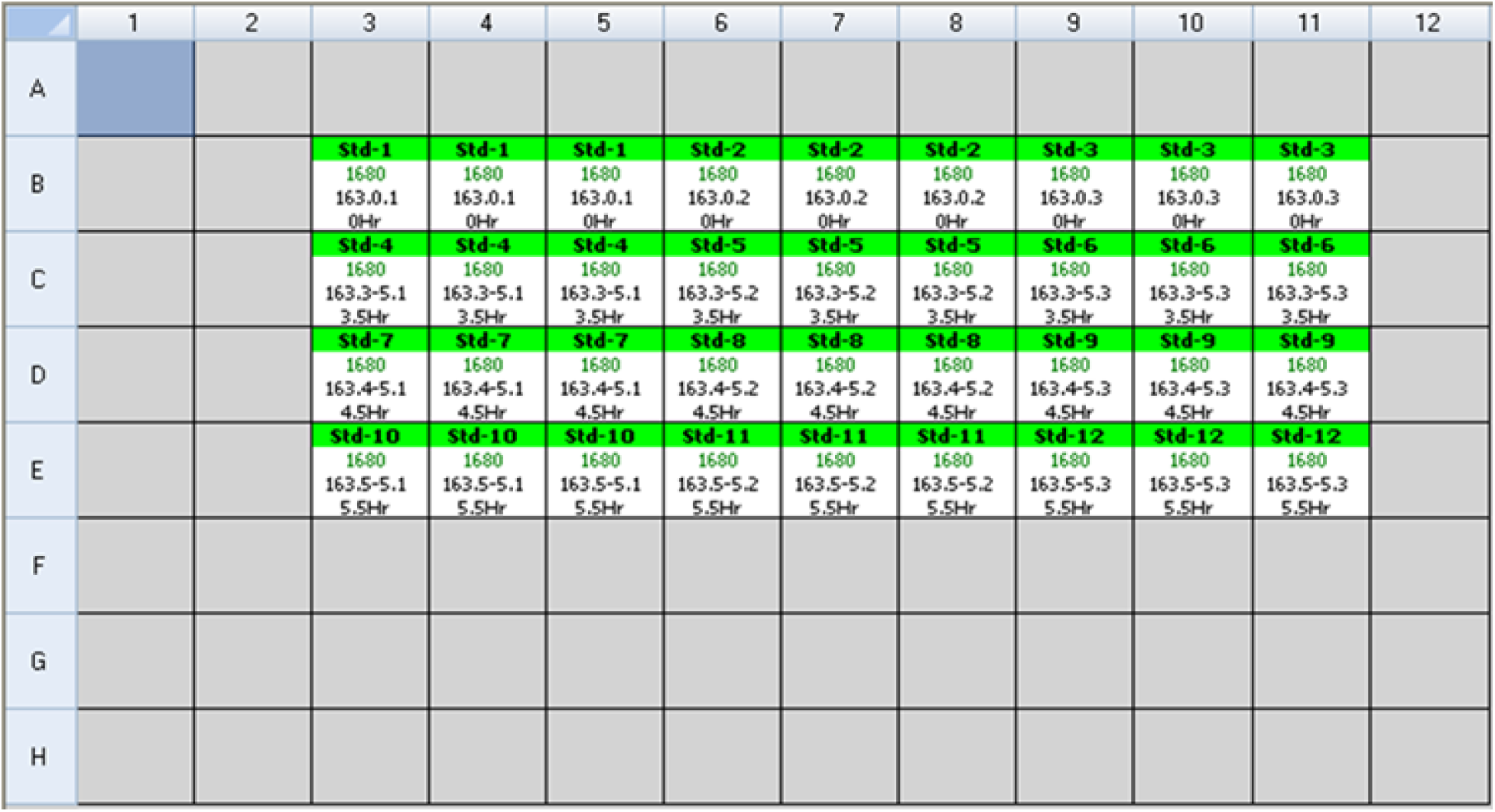
An example plate setup for one of the genes analyzed in the gene study. This is programmed via the BioRad CFX manager software.

### Data Analysis

Using the CFX manager software and the Gene Study menu, data was analyzed using Normalized Expression, or the ΔΔCq method. Experiment settings were defined so that T=0 time point was set as the control. Reference genes 1 and 3 were used to normalize the data (Reference gene 2 was deemed not a good reference gene). Efficiency and R^2^ values were entered into the experiment settings using data from standard curves. Biological Set Analysis was performed with Target vs. Time Point.

## Results

In order to analyze which candidate genes could be active conjointly with *GLR3.3* during the root gravitropic response, gene expression profiles for all candidate genes and reference genes were created. All primers were validated with standard curves (Table 2), with the exception of the third reference gene and 2180. The primer for 2180 could not produce an acceptable standard curve, even after primer re-design. Efficiency rates had to be within 95-105% according to MIQE guidelines; R^2^ had to be above 0.98 (Bustin 2009).

**Table 2:** Standard Curve Results.

| **Gene** | **Efficiency Value** | **R<sup>2</sup> Value** |
| --- | --- | --- |
| 3.3 (GLR 3.3) | 104.2% | 0.988 |
| 1680 (Myb Domain Protein) | 98.2% | 0.988 |
| 1840 (Potassium Channel) | 101.3% | 0.987 |
| 2180 (DC1-Domain Protein) | N/A | N/A |
| 2820 (PPR) | 95% | 0.982 |
| 10660 (Kinase) | 96.3% | 0.996 |
| Reference #2 | 104.9% | 0.986 |
| Reference #3 | - | - |

The three replicates show similar results (Figures 3a, 3b, 3c). *GLR3.3* expression remains relatively constant over time, with small changes, if any, compared to time zero. *GLR3.3* expression did increase by 50% in replicate 3 (Figure 3c). Two genes were observed to have expression patters that made them likely candidates as components of the *GLR3.3* pathway. *AtPK5* shows a distinct peak in expression at 4.5 hours in all three replicates with a slight decrease in expression at 5.5 hours. PPR shows an increase in expression with time in all three replicates (Figure 3).

**Figure 3:**
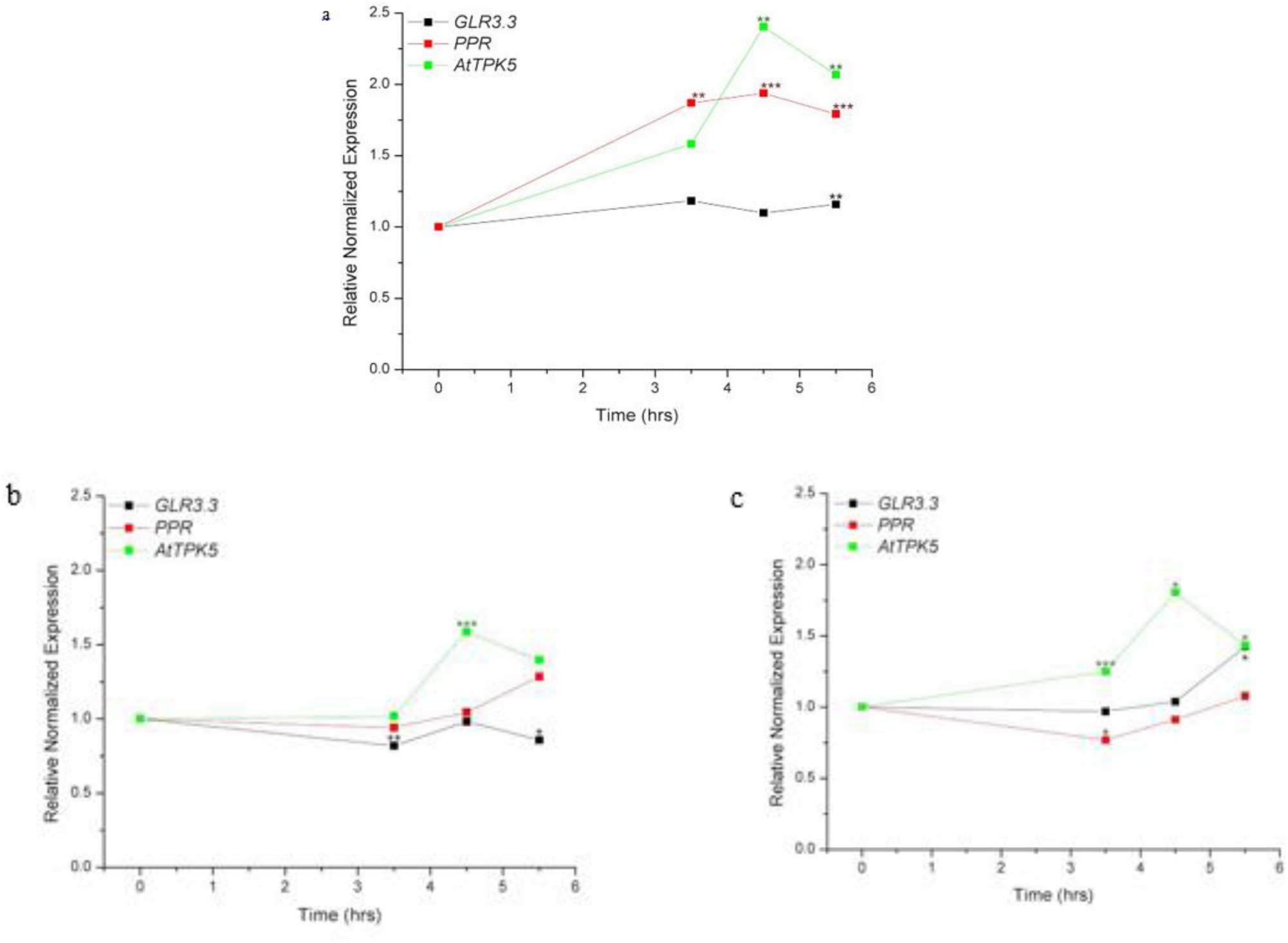
The relative normalized expression over time of *GLR 3.3* (At1G42540), Pentatricopeptide repeat-containing protein (PPR) (At4g02820), and AtTPK5 potassium channel (At4g01840) in three separate biological replicates: a, b, and c. Expression is compared to T=0: * = p < 0.1, ** = p < 0.5, *** = p < 0.01.

## Discussion

By identifying possible components of the gravitropic pathway that act concurrently with *GLR 3.3*, a novel metabolic pathway can begin to be described. Previous methods utilized to obtain qualitative data about gene expression through PCR (Merithew 2011) were not sensitive enough to accurately identify changes in gene expression. Using qRT-PCR, the initial six candidate genes were narrowed to two genes which are very likely part of GLR-dependent gravitropic cell signaling.

Of the six tested candidate genes, two show promising patterns that suggest involvement in the gravitropic cell signaling pathway. At4g01840 (AtPK5) shows a distinct pattern in all three replicates; gene expression peaks at the 4.5 hour time point and then decreases, with overall expression increasing over time (Figure 3). At4g02820 (PPR) shows a similar profile in all three replicates of overall increased expression (Figure 3). Additional published research supports our data that At4g01840, an AtTPK5 potassium channel, and At4g02820, a pentatricopeptide repeat-containing protein are likely components of this pathway, which is further discussed below.

Examining the *GLR3.3* expression profile (Figure 3), it is evident that gene expression remains fairly constant over time. This would be expected since this *GLR* is a membrane receptor, and would likely begin the gravitropic response. In a signal transduction pathway, the beginning components of the cascade would show less gene expression than components that are present in larger magnitudes toward the end of the signaling pathway.

Previous research has implicated At4g01840 (AtTPK5) as a cell signaling component. AtTPK5 channels are found in membranes of vacuoles in *A. thaliana* (Voelker 2006). These channels are voltage-independent potassium ion channels; thus, they could be activated by a ligand. When AtTPK5 is inhibited, the plasma membrane depolarizes easier, which increases cellular activity and thus increases responsiveness of the cell to other stimuli and calcium entry. This, in turn, can promote hormone release and downstream physiological responses. Voelker et al. speculate that because of these actions, *AtTPK* genes could function to fine tune the electrical properties of the membrane for specialized tasks. This idea fits well with the role of the GLR-dependent gravitropism pathway.

At4g02820 is a pentatricopeptide repeat protein (PPR). This family of proteins has a widely unknown role in plant physiology. Research shows that PPRs may be involved in organelle synthesis and bind to RNA to regulate gene expression (Narsai 2011). This suggests that PPR Proteins could be involved in gene regulation during gravitropism.

To further confirm that these genes are involved in the gravitropic response, they should be mutated and made non-functional. After this, seedlings should be studied in a way similar to Miller et. al. 2010. If seedlings with mutated At4g01840 and At4g2820 respond with a similar phenotype as seedlings with mutated *GLR3.3*, it would be highly likely that these genes are part of the gravitropic pathway. By utilizing qRT-PCR more accurate expression profiles were constructed, which were used to identify two genes that are likely components of the *GLR3.3* pathway. This is the start of a process to map the novel pathway of plant gravitropism.

## Appendix A

### Edit Protocol

Understanding the settings of the provided Bio-Rad CFX Manager software is important in obtaining accurate results. While some settings are available on the Thermocycler itself, it is easier to operate the instrument remotely from the connected computer. The plate set up was created before beginning any reactions. This process has to be done on the remote computer. This was done by opening the CFX manager software and selecting the “Run setup” and “User-defined”. The protocol editing screen will then appear. If a protocol has already been saved and created it can be found on the computer by using the “Select Existing. .”. This type of file will be a .prcl file. If it is necessary to create a new protocol the “Create New. .” button should be selected.

When creating a new protocol be sure to update sample volume input located at the top of the screen. To edit step temperatures or duration click directly on the value on either the graphical representation or the list below and type the new value once the value is selected. The buttons on the left hand side are how steps can be added to the protocol. To create a melt curve for primer select the “Insert Gradient” button. This will set the themocycler so that each row of the wells will run at a different temperature. When this button is selected it will show a gradient calculator on the right bottom corner. The best way to use the calculator is to insert the value for the average melting temperature that will be used for all primers into either the D or E value. The numbers that are shown on the gradient calculator directly correlate with the temperatures that the rows of the thermocycler will run at.

If the protocol requires a melt curve, it must be inserted after the repeating cycles of annealing. This is done by clicking the “Insert Melt Curve” button; temperature and duration values can be edited using the method previously mentioned. In this case a plate read will automatically be added to the melt curve. The most important thing to ensure when creating a protocol is to make sure that a plate read is added to the appropriate step, most likely the annealing/extension stage. It is often helpful to add a final holding step to the protocol. This is done by inserting a final step and changing the values to 4°C for a time of 0:00. This time will appear as “forever” when running the protocol.

### Plate Setup

The next tab in the “Run Setup” window of the software is dedicated to plate setup. Again, a previously designed plate can be selected from the computer. These will be saved as a .pltd file. To create a new plate select “Create New. .” and a clean slate will be provided. The “Scan Mode” setting on the top tool bar is defaulted to detect “All Channels”. To decrease the time of the instrument run, select “SYBR/FAM only” if using a Sybr Green enzyme. The Trace Styles option allows certain wells to be assigned specific colors, in order to help with initial visual analysis. When editing the plate, click or drag and click a group of wells that the setting will apply to. Sample Type must be chosen first. Sample name can be input into the box. This is typically the tissue used (i.e. 1/29/14 163 3.5). After typing it, check the “Load” box. Next, Indicate the Target gene.

To indicate technical replicates, use the replicate button by selecting all technical replicate wells and checking the load button. If all of the settings are not identical between the selected wells the replicate button will not work. To indicate a different set of technical replicates, arrow up so the Replicate # increases and then select the wells and hit load. A concentration of template should be chosen for each well. The concentration must be in scientific notation format.

Pay special attention to the orientation of the plate setup and ensure that you insert your PCR reaction plate in the correct direction. The lettering/numbering provided on a 96 well plate is helpful when attempting to orient the plate to the software. A diagram of the plate can be printed by right clicking the grid and selecting “print image”. This document is useful when preparing reactions away from the computer.

Once the protocol and plate setups have been selected or edited, progress to the “Start Run” screen. Here ensure the correct protocol and plate are correctly selected. It is possible to start the run and pause it just after the lid and block warm up. When this is done, the PCR reaction can proceed immediately. If it would be beneficial in a certain experiment to not have the PCR reaction to sit longer than normal, click start run. Immediately push the pause button when the option becomes available.

